# Neuroanatomy of catecholaminergic circuits in the brainstem and hypothalamus using T1-weighted and diffusion magnetic resonance imaging in humans: implications for brain-immune interactions, cardiovascular disease, neuropsychiatric disorders, stress response, and COVID-19

**DOI:** 10.1101/2025.04.11.644973

**Authors:** Poliana Hartung Toppa, Richard J. Rushmore, Kayley Haggerty, George Papadimitriou, Edward Yeterian, Agustin Castañeyra-Perdomo, Nikos Makris

## Abstract

Neuroimaging allows the study of brain structures that previously were undetectable due to their small size and location. Herein we focused on the core catecholaminergic circuitries in the human brain, involving the coerulean noradrenergic (or norepinephrine, NE) and dopaminergic (DA) systems. Using T1-weighted MRI morphometry and dMRI tractography, this study was carried out in one post-mortem human ultra-high-resolution dataset of the brainstem and diencephalon and in healthy human datasets from the Human Connectome Project repository. We investigated 26 connections of brainstem origin (13 in the left side and 13 in the right side) associated with the NE and DA circuitries. We delineated the coerulean NE and DA core central catecholaminergic circuitries of the brainstem and hypothalamus in the post-mortem dataset, including all targeted fiber connections. Importantly, this was also achieved in the HCP datasets. These results emphasize the importance of multispectral neuroimaging in the study of chemical neuroanatomical circuitries and its application in clinical conditions such as cardiovascular disease, major depression, schizophrenia, and other disorders associated with chronic stress and brain-immune interactions such as COVID-19.

## 1. Introduction

The three types of molecules contained in the human central nervous system (CNS) and referred to as catecholamines (or central catecholamines) are norepinephrine (NE) (or noradrenaline), epinephrine (E) (or adrenaline) and dopamine (DA). These molecules share a common biochemical origin. They originate from the L-amino acid tyrosine, which by a series of successive enzymatically-operated conversions gives rise to NE, E and DA. The topographical and structural organization of the catecholamine-containing neurons shows a notable consistency in the mammalian brain. These cell groups are located principally within the brainstem and the hypothalamus. In particular, NE and E are produced in the pons and medulla, whereas DA is produced in the mesencephalon (midbrain) and the hypothalamus (1–3). The neurochemistry and structural connectivity of the catecholamines have been studied extensively since the 1960s in experimental animals and, more recently, in humans. This research has resulted in the integration of chemical neuroanatomy with traditional neuroanatomy.

The biobehavioral and clinical effects of the catecholamines are numerous and remarkable. NE and E play a key role in cardiovascular and respiratory regulation and in stress response, motivation, mood regulation and brain-immune interactions. Dopamine is central in reward function, motivation and mood regulation. Moreover, it plays a role in cognitive processing including memory and attention as well as in many other bodily functions, especially movement. Thus, the understanding of catecholaminergic human brain circuits is key in basic and clinical neuroscience. Furthermore, given the nature of clinical practice, which necessitates both molecular-metabolic and anatomical-histological knowledge and clarity associated with therapeutic interventions, the identification of these systems and the ability to monitor their structure and function in health and disease is a matter of great clinical relevance. In this regard, the advent of neuroimaging has been of great advantage, given its non-invasive and in vivo nature that has resulted in the development of novel research and clinical opportunities. The catecholaminergic systems in the brainstem and hypothalamus comprise specific cells of origin and axonal fiber pathways, which course via specific topographic trajectories toward their terminations (2,3). NE circuits, in particular, can be distinguished as coerulean NE, which are associated with the locus coeruleus (LC), and non-coerulean NE, which are associated with NE cell groups within the brainstem that are not located in the LC.

Given the complex anatomy of the brainstem and the hypothalamus as well as the spatial limitations these brain regions present for neuroimaging, clinical structural neuroimaging investigations of fine-grained brainstem and hypothalamic anatomy are extremely challenging (4). In this regard, multispectral neuroimaging allows the identification of brain gray matter structures as well as the fiber tracts that interconnect them. More specifically, T1-weighted magnetic resonance imaging (MRI) morphometry enables the segmentation of brainstem, diencephalic and other cerebral gray matter regions of interest (ROIs), whereas diffusion MRI (dMRI) tractography permits the delineation of white matter fiber pathways. Thus, the brain circuits associated with coerulean NE and DA systems, which are constituted by gray matter cortical regions, subcortical nuclei and their associated axonal white matter fibers, can be sampled and measured reliably using current neuroimaging.

In the present study we report the results of the coerulean NE circuitry in 12 Human Connectome Project (HCP) subjects. Five of these subjects were analyzed in a previous pilot publication by our group (5). In this study we analyzed seven additional HCP subjects, thus creating a preliminary database of the coerulean NE circuitry in a balanced cohort of six females and six males matched in age from the HCP repository. Furthermore, we report here for the first time preliminary observations on the human dopaminergic structural circuitry involving regions within the brainstem and hypothalamus. To this end, we first delineated the DA circuitries in one ultra-high-resolution post-mortem human dataset (4). This was done using combined structural T1-weighted MRI morphometric and dMRI tractographic analyses. We then used the same combined method to delineate the DA circuitries in five healthy human subjects from the publicly available HCP repository (5). Importantly, we reconstructed the mesostriatal, mesolimbic and mesocortical DA projection systems in three-dimensional space. The five subjects used to delineate the DA circuitries have been used in a recent study by our group (6) to delineate the coerulean norepinephrine circuitry. Thus, we complemented our prior study of the dorsal catecholaminergic systems by investigating the more ventrally located DA systems in the brainstem and hypothalamus. Moreover, we measured biophysical parameters of average fractional anisotropy (FA) (7), axial diffusivity (AD) (8) and radial diffusivity (RD) (8,9) of all fiber tracts in the 12 healthy human subjects used to delineate the coerulean NE circuitry and the DA circuitry in the five healthy human subjects. Thus, we produced comprehensive visualizations of the central catecholaminergic structural circuitries, both coerulean NE and DA, in the human brain.

## 2. Materials and Methods

### 2.1. Rationale

We studied the structural connectivity of central catecholaminergic systems, specifically coerulean NE and DA circuitries following the anatomical literature, which we have elaborated on in detail under “Structural anatomy and topographical organization of catecholamine systems” in the discussion. There were 26 connections (13 in the right side of the brain and 13 in the left) studied herein. Two connections involved NE circuitry, namely NST/DMN (nucleus of the solitary tract/dorsal motor nucleus of the vagus) – LC (locus coeruleus) and LC – PVN (paraventricular nucleus)/Hypothalamus. Eleven connections involved DA circuitry, as follows: SN (substantia nigra) – Thalamus, SN – Putamen, SN – Entorhinal, SN – Superior Frontal, SN – Caudate, SN – Cingulate, SN – Orbitofrontal, VTA (ventral tegmental area) – Thalamus, VTA – Nucleus Accumbens, VTA – Cingulate, and VTA – Orbitofrontal. To determine these circuits using MRI, we followed the multispectral neuroimaging approach of “MRI-based brain volumetrics” (10), a morphometric neuroscience framework developed at the Center for Morphometric Analysis (CMA) at Massachusetts General Hospital/Harvard Medical School and the A. A. Martinos Center for Biomedical Imaging since the 1990s (10–16). This framework gave rise to the Harvard Oxford Atlas (11) as distributed in FSL in the early 2000s and served as the testbed for validation of the most currently available automated methods for brain morphometry, in particular FreeSurfer (17,18). In this approach, the definition of brain ROIs is based on landmarks of the individual brain under investigation such as sulci, protuberances or bulges and gray-white matter contrasts, which are reliably identifiable in different neuroimaging modalities (14,19,20). This approach requires a thorough knowledge of neuroanatomy, which is critical for the study of structures with complex neuroanatomical morphology and architecture such as the brainstem and hypothalamus (12,14). Currently available techniques and protocols generally used for clinical neuroimaging do not have the necessary spatial resolution enabling the visualization of small structures such as nuclei. Thus, to reliably localize the brainstem and hypothalamic nuclei, it is necessary to follow conventions with respect to anatomic morphologic landmarks that are consistently identifiable using lower spatial resolution MRI datasets and compare them with histological atlases or ultra-high-resolution datasets such as that of Calabrese and colleagues (2020) (4,14) in which these anatomic landmarks are also identifiable. This approach has been used traditionally in anatomic radiological and anatomic clinical studies and was also adopted by neuroimaging morphometric analysis since the early 1980s (10,21). In **Table 1** we show the demographics of the HCP subjects analyzed for the coerulean noradrenergic (NE) and dopaminergic (DA) circuitries. We investigated the coerulean circuits in 12 healthy HCP subjects. Note that the five HCP subjects in which we delineated the DA systems are the same subjects we analyzed in a prior study by our group of the coerulean NE circuits (6). By expanding our prior analysis of coerulean NE circuits (6) from five subjects to 12 in the present study, we initiated a preliminary database of these circuits in humans. We investigated the DA systems in the brainstem and hypothalamus as follows. We first delineated the DA circuitries in one ultra-high-resolution post-mortem human dataset (4) and then used the same method to delineate the DA circuitries in five healthy human subjects from the HCP repository (5). By using an overlapping sample of five subjects for both the coerulean NE and DA circuitries, we aimed to distinguish the more dorsally located coerulean NE circuits from the more ventrally located DA circuits in these same five individual HCP brains.

**Table 1.** Demographics of Human Connectome Project repository subjects analyzed in this study. The subject identification numbers (IDs), female (F) or male (M) gender, and ages (in years) are specified per the HCP database. (Abbreviations: DA, dopamine; NE, norepinephrine)

| Subject ID | Gender | Age | DA | NE |
| --- | --- | --- | --- | --- |
| 100307 | F | 26-30 | X | X |
| 101107 | M | 22-25 | X | X |
| 101915 | F | 31-35 | X | X |
| 103111 | M | 26-30 | X | X |
| 103414 | F | 22-25 |  | X |
| 110411 | M | 31-35 | X | X |
| 111312 | F | 31-35 |  | X |
| 114419 | M | 31-35 |  | X |
| 118528 | F | 26-30 |  | X |
| 118932 | M | 26-30 |  | X |
| 124422 | F | 31-35 |  | X |
| 129028 | M | 26-30 |  | X |

### 2.2. Morphometric delineation of the DA and coerulean NE regions of interest (ROIs) and dMRI tractographic analysis in the ultra-high-resolution post-mortem dataset

The morphometric approach for the delineation of the DA and coerulean NE circuits involved two procedural steps. In Step 1 we segmented the relevant structural ROIs associated with the circuits in the brainstem and hypothalamus. We also segmented the thalamus as a whole. More specifically, using the b0 dMRI structural dataset we segmented the ROIs where the cells of origin of the DA (**Figure 1A**) and LC-related (**Figure 1B**) central catecholamine systems are located. DA cells are located in the midbrain, namely the substantia nigra including cell group A9/substantia nigra pars compacta (SNpc) and the A10/ventral tegmental area (VTA) (22–24). In the diencephalon, DA neurons of origin are located mainly in the hypothalamus. We labeled the ROIs associated with the DA cells of origin as substantia nigra (SN), VTA and PVN as shown in **Figure 1A**. We also labeled the structures associated with the coerulean norepinephrine (NE) system, namely the locus coeruleus (LC) in the upper pons, the nucleus tractus solitarius (NTS) and the dorsal motor nucleus of the vagus (DMN) in the upper medulla (B1) and the paraventricular nucleus (PVN) in the hypothalamic region of interest (ROI) as shown in **Figure 1B**. In the present study we subdivided further the dorsal vagal complex (DVC) into nucleus tractus solitarius (NTS) and the dorsal motor nucleus of the vagus (DMN), which we did not do in our prior study (6) in which DVC was considered as a whole. Given the differential connectivity of the LC with the NTS and DMN, this complex circuitry involving the LC could play an substantial role in brain-immune interactions (25,26). In Step 2 the ROIs identified during Step 1 were used as “seeds” for dMRI tractography, which was performed to delineate the DA structural connections. To obtain the different fiber tracts that reached the a priori selected cerebral cortical regions, we used the White Matter Query Language (WMQL, (27)). Note that the WMQL was not used in our prior study of coerulean NE circuitry (6).

**Figure 1.**
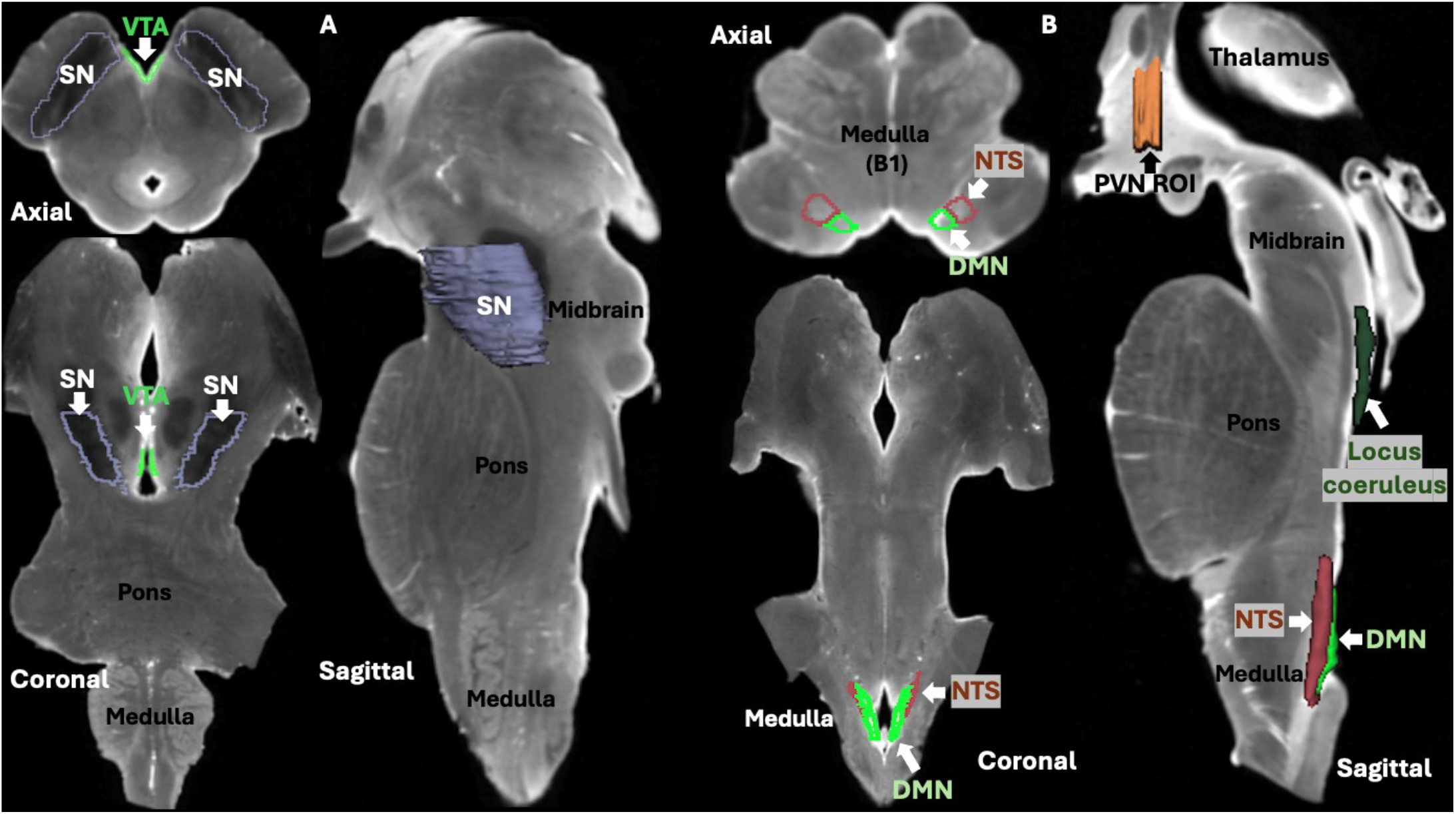
Method used in the human ultra-high-resolution post-mortem dataset for structural MRI-based delineation of the cell groups or nuclei that give origin to the dopaminergic (DA) and coerulean norepinephrine (NE) central catecholamine circuitries. **Figure 1A** shows the structures associated with the DA system, namely the substantia nigra (SN) and the ventral tegmental area (VTA) in the midbrain. **Figure 1B** shows the structures associated with the coerulean nore-pinephrine (NE) system, namely the locus coeruleus (LC) in the upper pons, the nucleus tractus solitarius (NTS) and the dorsal motor nucleus of the vagus (DMN) in the upper medulla (B1) and the paraventricular nucleus (PVN) in the hypothalamic region of interest (ROI).

Currently, diffusion MRI data at 200 micron isotropic voxel resolution, such as the Calabrese et al. (4) post-mortem dataset used in this study, can be considered as ground truth in the field of diffusion MRI. Therefore, the delineation of catecholaminergic circuits in our ultra-high-resolution post-mortem human dataset may serve as a ground truth comparator for the commonly used HCP datasets. That said, it must be emphasized that validation of human connectional neuroanatomy draws heavily upon animal experimental findings, given the lack of tracing techniques in humans comparable to those in nonhuman species and the known limitations of dMRI tractography in humans. This notion has been elaborated upon in recent publications by our group (13,15). Moreover, it is critically important to realize that structural connectivity of the brainstem and the diencephalon including the hypothalamus as well as the limbic system, based on experimental research on non-primate mammals (e.g., rodents, cats, dogs), is highly comparable to humans. However, this is not the case for the neocortex, which is considerably increased in size and complexity in primates, especially humans. Thus, our analysis of the present neuroimaging data, which is largely focused on brainstem, diencephalic and limbic NE and DA circuits, has been carried out in the context of the known connectivity of catecholaminergic systems in non-primate mammals and nonhuman primates, which represent the current gold standard in connectional neuroanatomy of these brain circuits (see, e.g., 1,3).

### 2.3. Morphometric delineation of the DA and coerulean NE regions of interest (ROIs) and dMRI tractographic analysis in healthy HCP subjects

The morphometric approach for the delineation of the DA and coerulean NE circuits in healthy HCP subjects also involved two procedural steps. In Step 1 we segmented the ROIs that contain the cell groups of origin of the dopaminergic (DA) and coerulean noradrenergic (NE) central catecholamine circuitries. Specifically, we segmented the DA cell groups of origin, namely the substantia nigra (SN) and ventral tegmental area (VTA), and the coerulean NE cell groups of origin as shown in **Figure 2**. The coerulean NE neurons of origin are located in the pons. Herein, we sampled the LC with its neighboring pontine region. The pontine coerulean NE complex is constituted principally by cell group A6/LC and by adjoining noradrenergic neurons in the central gray and subcoerulean regions. Collectively, these cell groups comprise approximately half of the central NE brain system. We have reported on the LC NE cell groups in a recent publication (6) in five healthy HCP subjects. In the present study, we extended this approach to a larger population of 12 HCP subjects to generate a preliminary normative database for the human coerulean NE circuitry. Segmentation of the anatomical ROIs or parcellation units (PUs) was executed with the neurosegmentation module of the publicly available 3D Slicer software platform (28). In Step 2, the segmented anatomical PUs were sampled and used as “seeds” for tractography. In this step, dMRI-based tractography was performed for the delineation of the DA and coerulean NE structural connectivity. The mapping of these fiber tracts was done in individual subject space. To obtain the LC connections that reached the a priori selected cerebral cortical regions, we used the White Matter Query Language (WMQL, 27) rather than the 3D Slicer platform used in our prior study of this connectivity (6).

**Figure 2.**
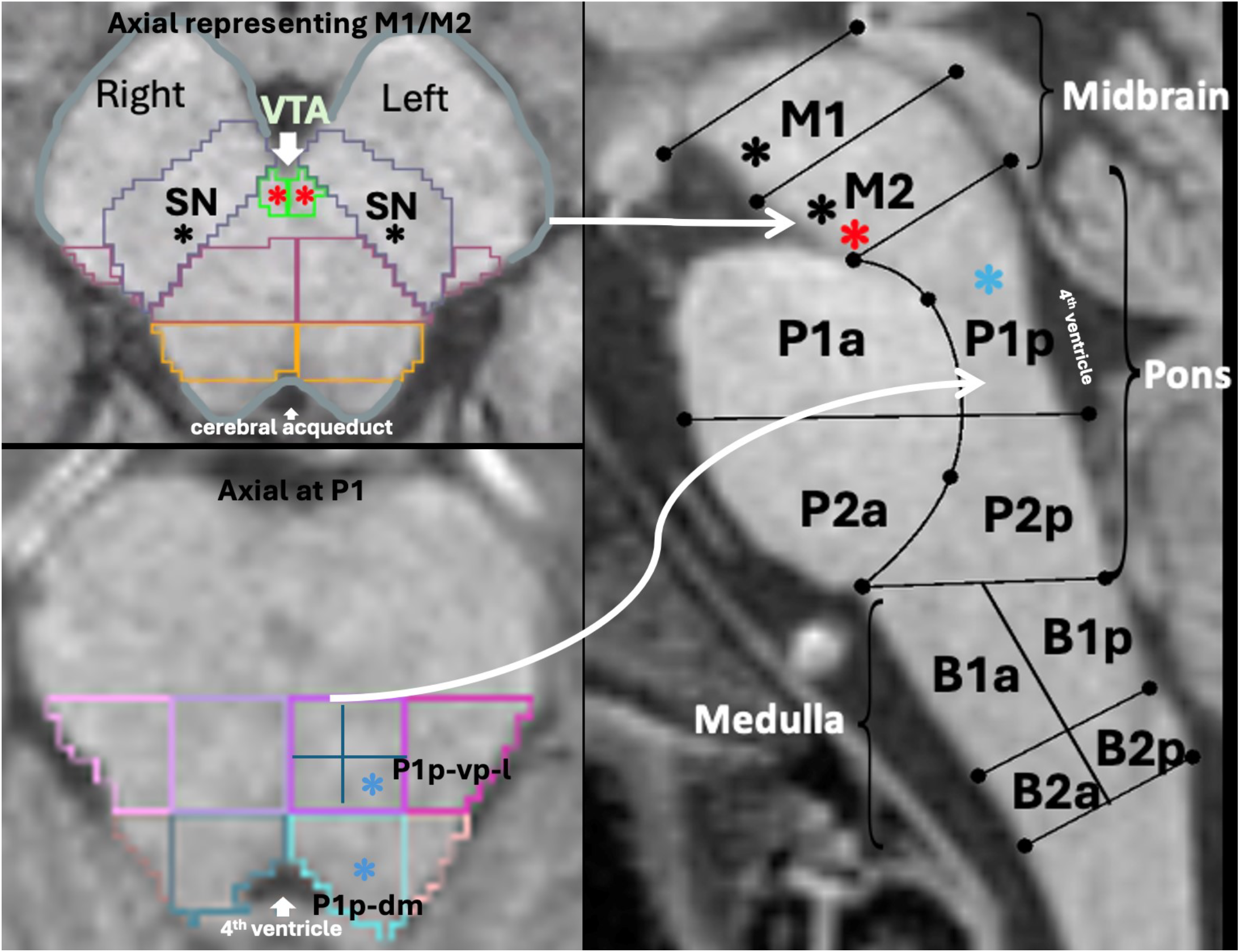
Method for structural MRI-based delineation of the regions of interest (ROIs), which contain the cell groups of origin of the dopaminergic (DA) and coerulean norepinephrine (NE) central catecholamine circuitries in one representative HCP in vivo dataset. The midsagittal section of the brainstem shows how morphometric parcellation was performed on T1-weighted images for the entire brainstem, which has been described in detail by DaSilva et al. (29) and Makris and colleagues (6). Using this procedure, we subdivided the brainstem into the midbrain (M1 [upper midbrain], M2 [lower midbrain]), anterior and posterior pons (P1a and P1p [upper pons]; P2a and P2p [lower pons]), and anterior and posterior medulla (B1a and B1p [upper medulla]; B2a and B2p [lower medulla]). The DA cell groups of origin, namely the substantia nigra (SN, indicated by black asterisks) and ventral tegmental area (VTA, indicated by red asterisks), are shown in the upper axial section of the left panel at the midbrain level in both sides of the brainstem. The coerulean NE cell groups of origin shown in the lower axial section of the left panel are contained in parcellation units (PUs) P1p_dm (upper pons posterior dorsomedial) and P1p-vp-l (upper pons posterior ventral posterior lateral) and are indicated by blue asterisks.

### 2.4. Neuroimaging protocols and dMRI tractographic analysis in the ultra-high-resolution post-mortem dataset

The human post-mortem brainstem and diencephalon dataset used in this study has been made available to the imaging research community and described in detail by Calabrese and colleagues (2015) (4). Briefly, imaging was performed in a single human brainstem and diencephalon from an anonymous 65-year old male donor as described by Calabrese et al. (2015), following our institutional rules and guidelines. The brain was extracted with a post-mortem interval of 24 h and the brainstem and diencephalon were dissected. The vasculature was cleared with normal heparinized saline (100 UI/ml heparin) and then the brain was immersed in 10% neutral buffered formalin for 2 weeks. A week prior to scanning, the sample was immersed in a 0.1 M phosphate buffer containing 1% gadoteridol. Prior to imaging, the brain was placed in liquid fluorocarbon (Galden PFPE, Solvay Plastics, Brussels, Belgium). RF transmission and reception were carried out with a 65 mm inner-diameter quadrature RF coil. With respect to the diffusion MRI protocol, Calabrese and colleagues provide the following description (2015) (4): “Postmortem MR imaging was performed in a 7 Tesla small animal MRI system controlled with an Agilent console (Agilent Technologies, Santa Clara, CA). Diffusion data were acquired using a simple diffusion-weighted spin echo pulse sequence (TR 5 100 ms, TE533.6 ms, BW5278 Hz/pixel). Diffusion preparation was achieved with a pair of unipolar, half sine diffusion gradient waveforms of width (d) 5 4.7 ms, separation (D) 5 26 ms, and gradient amplitude (G) 5 50.1 G/cm. Single-shell high angular resolution diffusion imaging (HARDI) data were acquired with 120 unique diffusion directions at b 5 4,000 s/mm2 and 11 b 5 0 s/mm2 (b0) volumes dispersed evenly throughout the acquisition. The FOV was 90 3 55 3 45 mm, and the acquisition matrix was 450 3 275 3 225 resulting in a 200 mm isotropic voxel size. Total acquisition time was 208 h.” For the tractographic reconstruction we used the White Matter Query Language (WMQL, (27)). Note that the WMQL was not used in our prior study of coerulean NE circuitry (6).

### 2.5. Neuroimaging protocols and dMRI tractographic analysis in healthy HCP subjects

Datasets were acquired from the HCP, as described in [HCP] (https://www.humanconnectome.org). ACPC-aligned T1w MRI images (0.7 × 0.7 × 0.7 mm^3^ voxel size) from the HCP Young Adult dataset were used for structural MRI analysis. Diffusion MRI analysis was performed on the same subjects. The T1-weighted and dMRI protocols are as follows. T1w: 3D MPRAGE, TR=2400 ms, TE= 2.14 ms, TI= 1000 ms, Flip angle 8 deg, voxel size 0.7 mm isotropic. Diffusion MRI: Spin-echo EPI, TR=5520 ms, TE=89.5ms, flip angle 78 deg, refocusing flip angle: 160 deg, multi factor 3, Echo spacing 0.78 ms, voxel size 1.25 mm isotropic, b values 1000, 2000 and 3000 s/mm^2^, each b shell with 90 diffusion directions and 6 b=0s. Whole brain tractography was performed using a multi-tensor unscented Kalman filter (UKF) method (20,22) as implemented in 3D Slicer. This algorithm uses tractography to drive the local fiber model estimation, i.e., model estimation (in this case, the multiple tensors) is done while tracing a “fiber” from seed to termination (23).

### 2.6. Quantitative Analysis

Biophysical parameters of average fractional anisotropy (FA), axial diffusivity (AD) and radial diffusivity (RD) of the fiber tracts in 12 healthy human subjects, 5 for the DA circuitry and 12 for the coerulean NE circuitry, were measured. The coerulean NE cell groups of origin shown in the lower axial section of the left panel in **Figure 2** are contained in parcellation units (PUs) P1p_dm (upper pons posterior dorsomedial) and P1p-vp-l (upper pons posterior ventral posterior lateral) and are indicated by blue asterisks. Inter-rater reliability of the ROIs in the upper medulla for the dorsal vagal complex (DVC) “seeding” and the upper pons for LC “seeding” was excellent as reported in a prior study by our group (4). The Dice coefficient was 0.99 for intra-rater reliability and 0.95 for inter-rater reliability. The presence or absence of specific fiber tracts in the left or right hemisphere of each individual subject was also recorded.

## 3. Results

We were able to delineate the DA circuits in one ultra-high-resolution post-mortem human dataset of the brainstem and diencephalon (**Figure 3**) as well as in five healthy human subjects of the HCP repository (**Figure 4**). Furthermore, we reconstructed the three principal dopaminergic projections, namely the mesostriatal (Figure 6), mesolimbic (**Figure 7**) and mesocortical (**Figure 8**). With respect to the coerulean NE circuitry we delineated the connections in the ultra-high-resolution post-mortem human dataset of the brainstem and diencephalon (**Figure 5**) as well as in 12 healthy human subjects of the HCP repository (**Figure 5**). In the present study, we expanded our observations of the human coerulean NE circuitry from five subjects originally reported by our group in a prior publication (6) to 12 subjects balanced for age and sex. Although small in size, it provides the first available preliminary normative database of the coerulean NE circuitry in humans.

**Figure 3.**
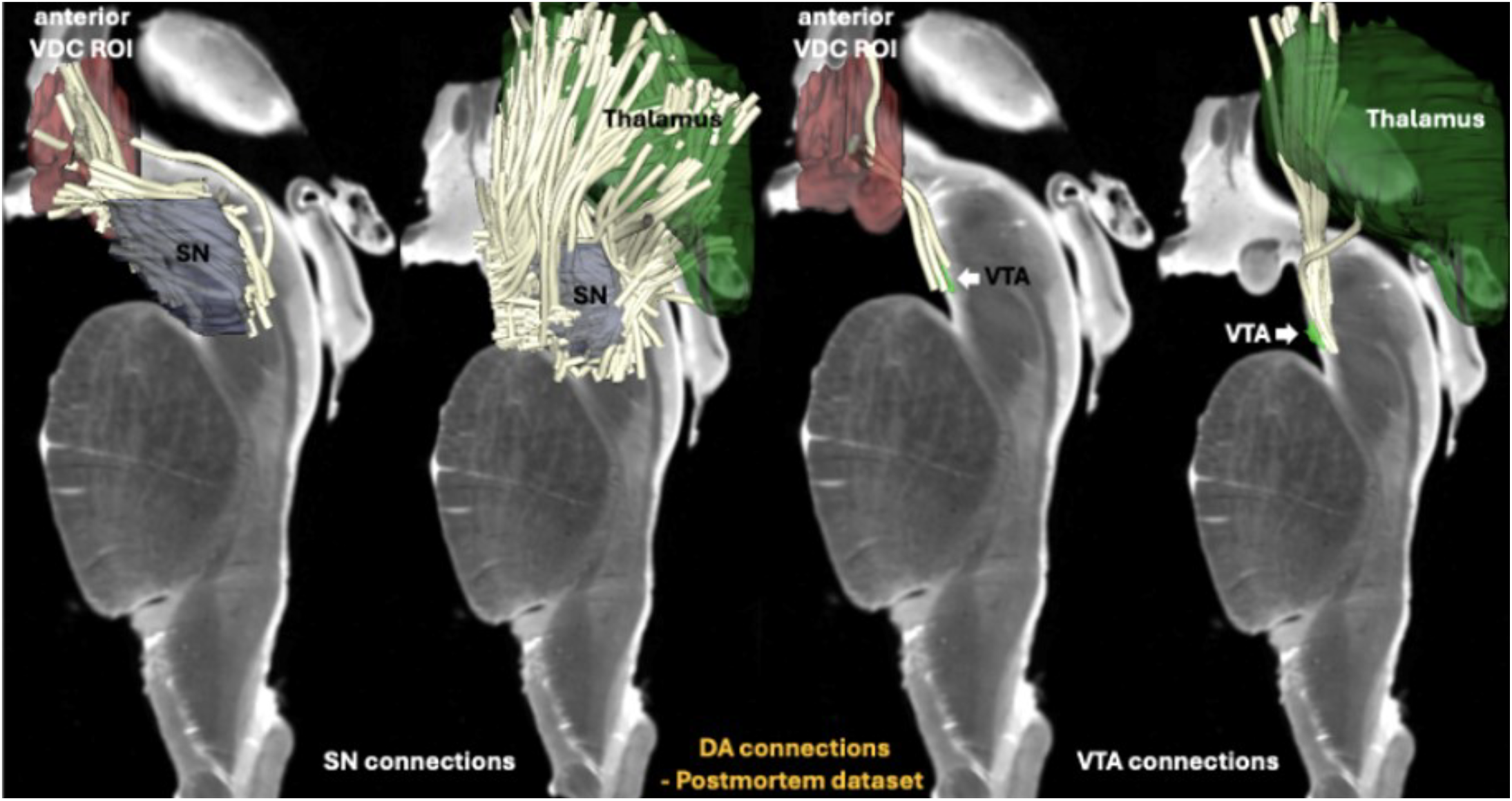
The results of dMRI tractographic analysis for the human ultra-high-resolution post-mortem dataset are shown for the left side of the brainstem and diencephalon. The dopaminergic (DA) connections of the substantia nigra (SN) and ventral tegmental area (VTA) with the thalamus and the anterior ventral diencephalic (anterior VDC) region of interest (ROI) are depicted in the four sagittal representations. It should be noted that the SN could not be subdivided into pars compacta and pars reticulata, thus it was sampled as a single ROI including both parts.

**Figure 4.**
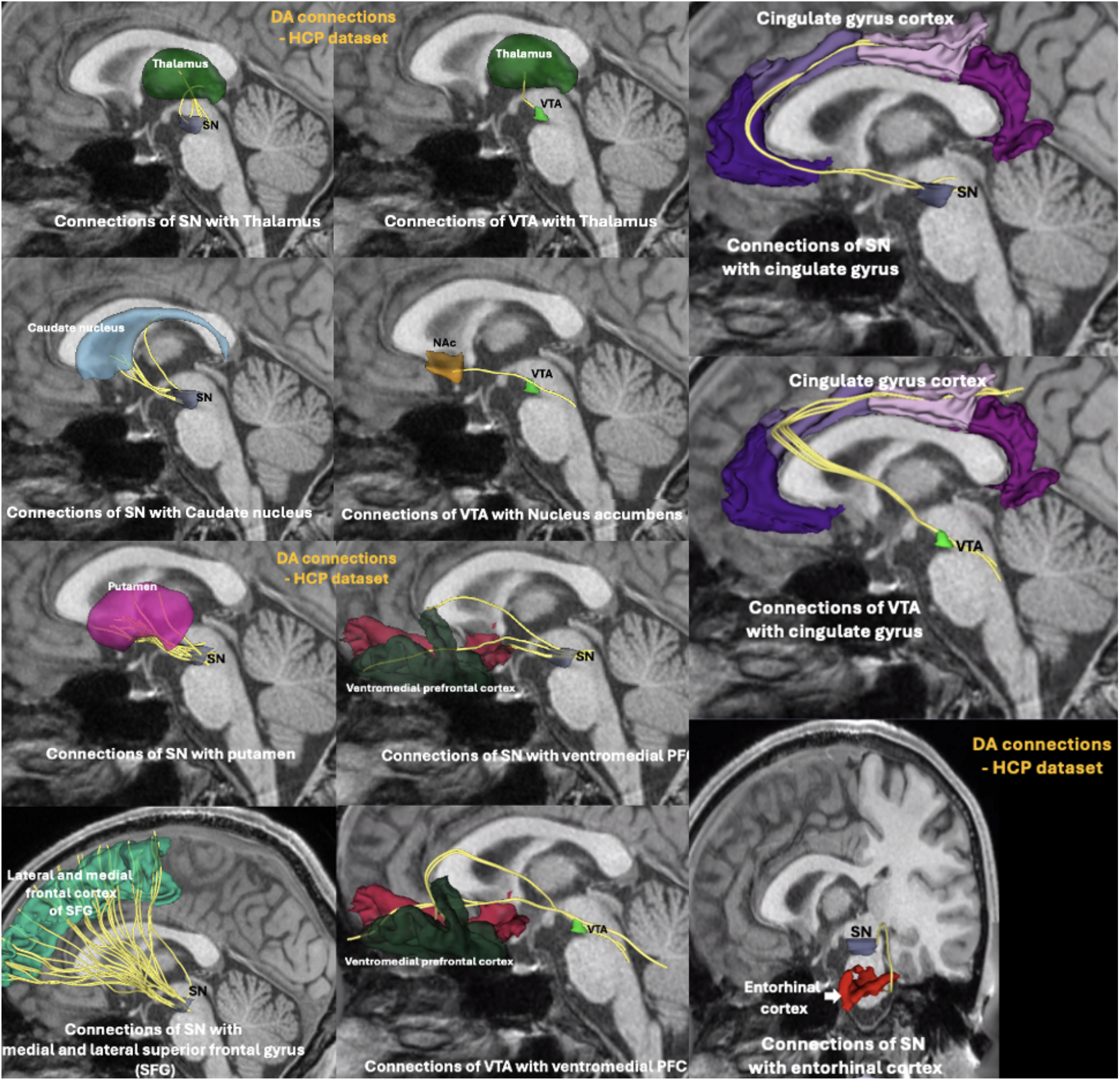
The results of dMRI tractographic analysis for Subject “HCP_110411” are shown for the left side of the brain. The eleven panels in the figure show the dopaminergic (DA) connections of the substantia nigra (SN) and ventral tegmental area (VTA) with the thalamus, caudate nucleus, nucleus accumbens (NAc), putamen, cingulate gyrus, ventro-medial prefrontal cortex (including the orbitofrontal cortex and medial prefrontal cortex) and the entorhinal cortex. It should be noted that we were not able to subdivide the SN into pars compacta and pars reticulata, thus we sampled SN as a single ROI.

**Figure 5.**
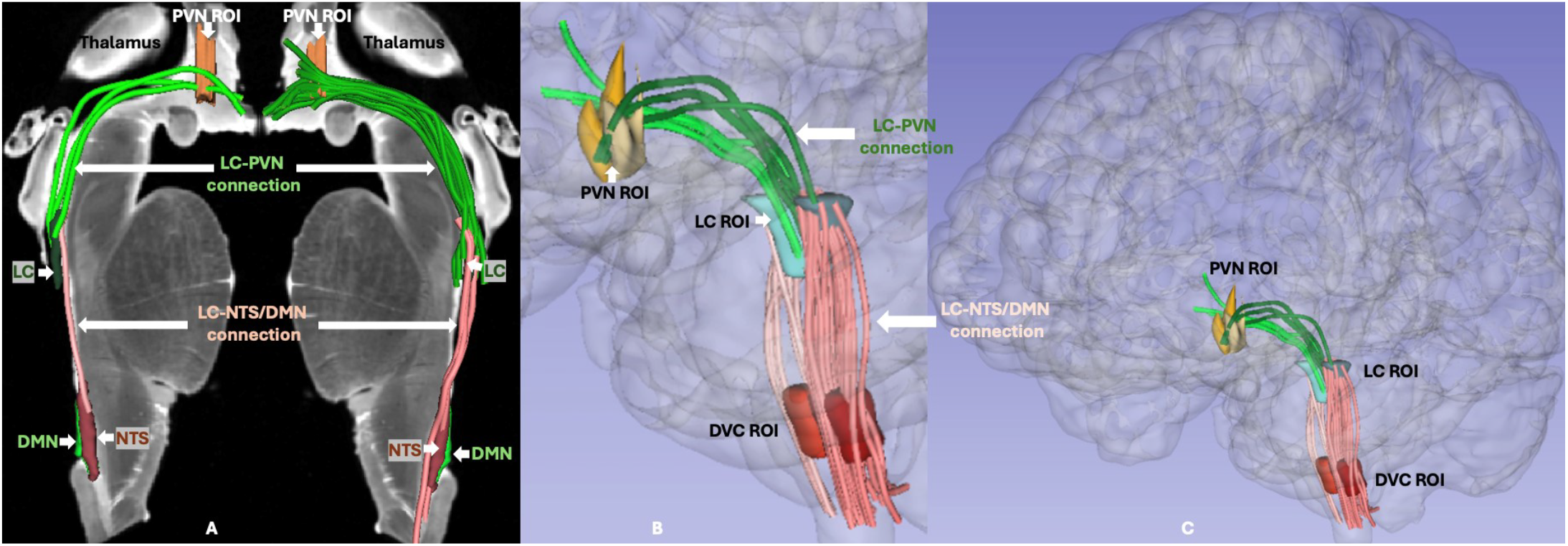
The coerulean norepinephrine (NE) cell group connections in human post-mortem and in vivo neuroimaging datasets. Panel A shows the results of dMRI tractographic analysis for the human ultra-high-resolution post-mortem dataset in the left and right side of the brainstem and diencephalon. Likewise, the results of dMRI tractographic analysis for Subject “HCP_110411” are shown in panels B (magnified detail of panel C) and C. Panels B and C show the LC circuitry within a “glass brain” visualization.

**Figure 6.**
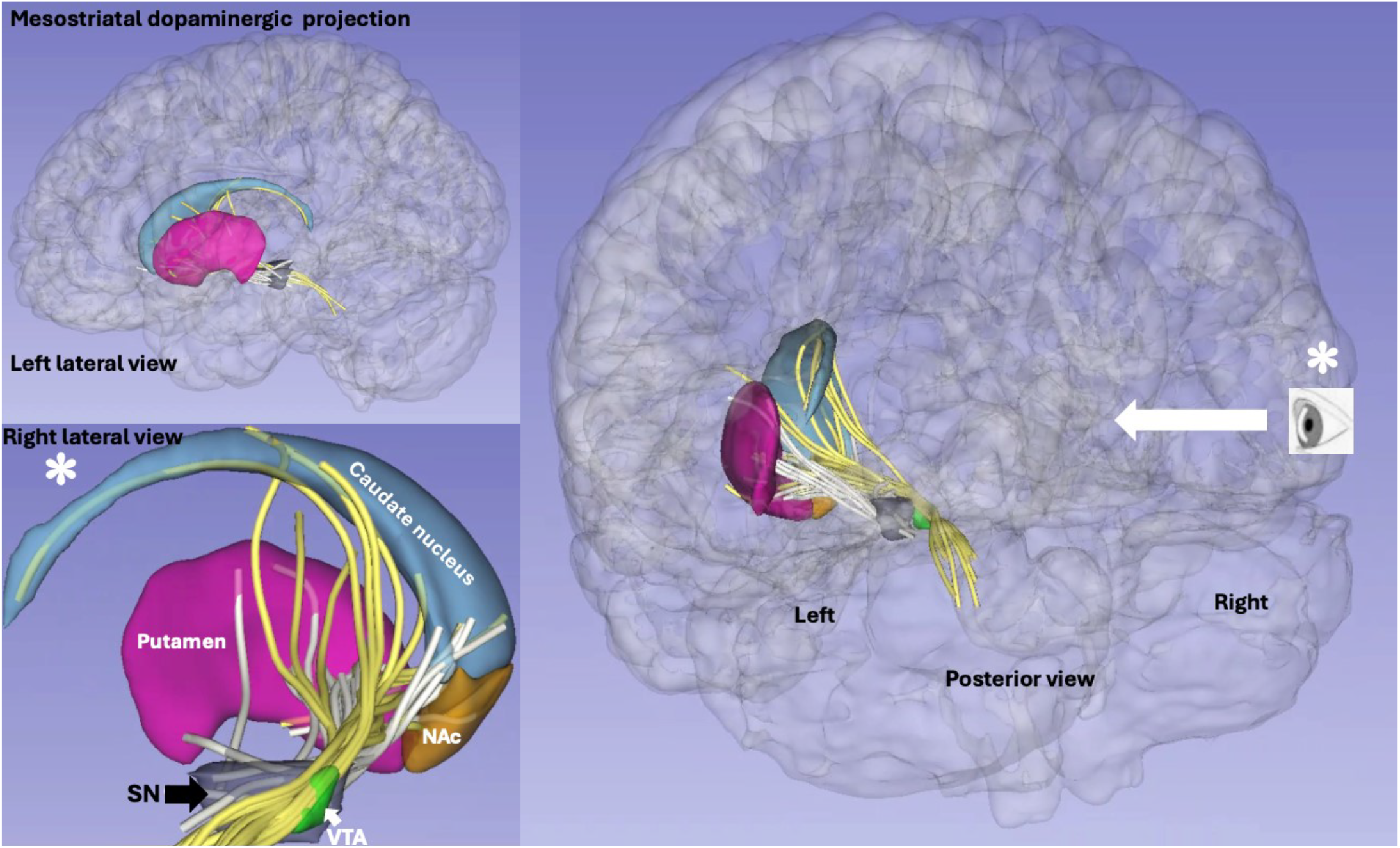
Mesostriatal dopaminergic (DA) projection system involving projections from the substantia nigra (SN) and the ventral tegmental area (VTA) to the caudate nucleus, putamen and nucleus accumbens (NAc). White asterisk denotes that the scene in the left lower panel should be viewed from a visual angle indicated by the white arrow in the right panel.

**Figure 7.**
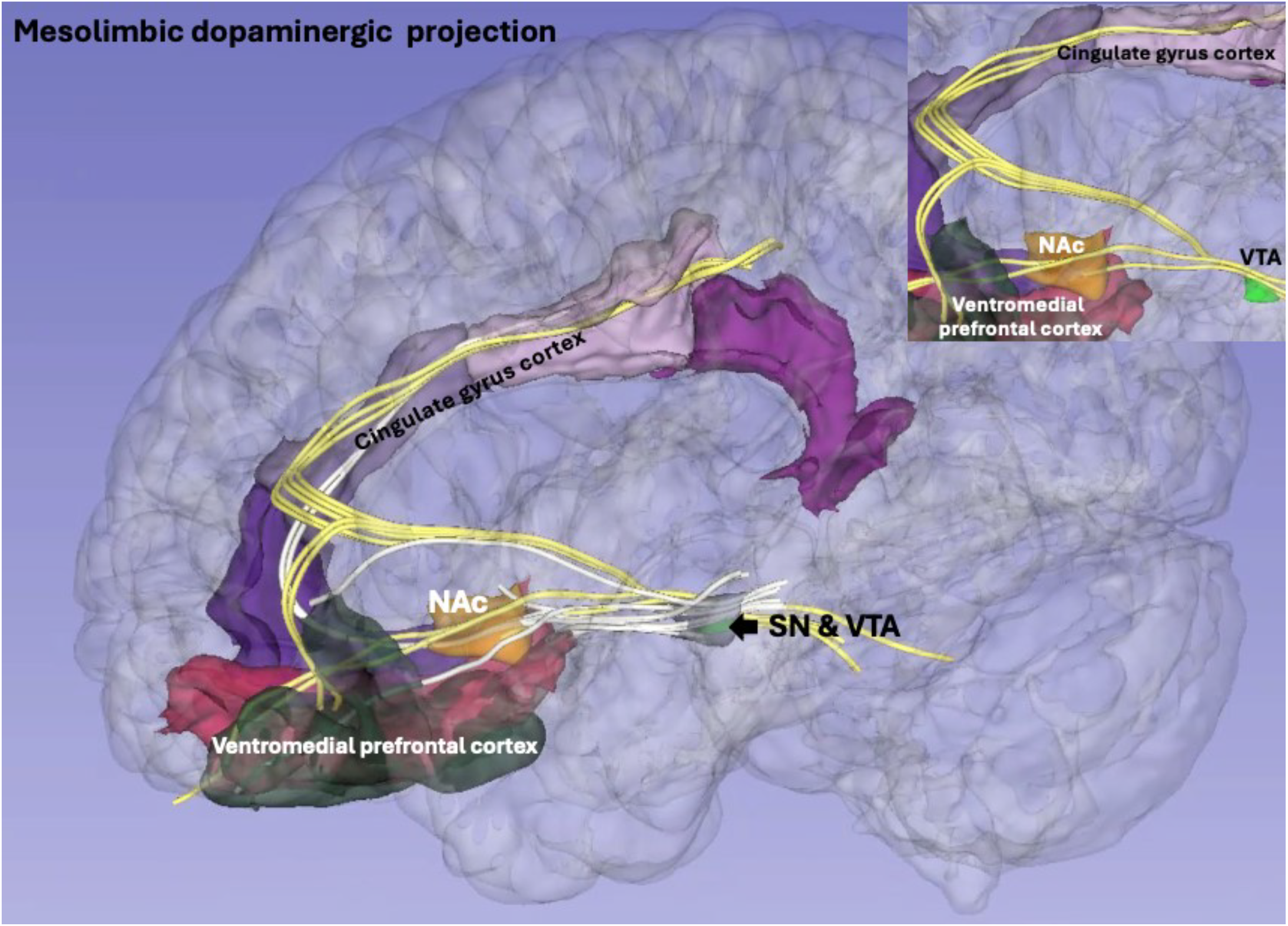
Mesolimbic dopaminergic (DA) projection system involving connections principally from the ventral tegmental area (VTA) and the substantia nigra (SN) to the nucleus accumbens (NAc), cingulate gyrus and ventromedial prefrontal cortex.

**Figure 8.**
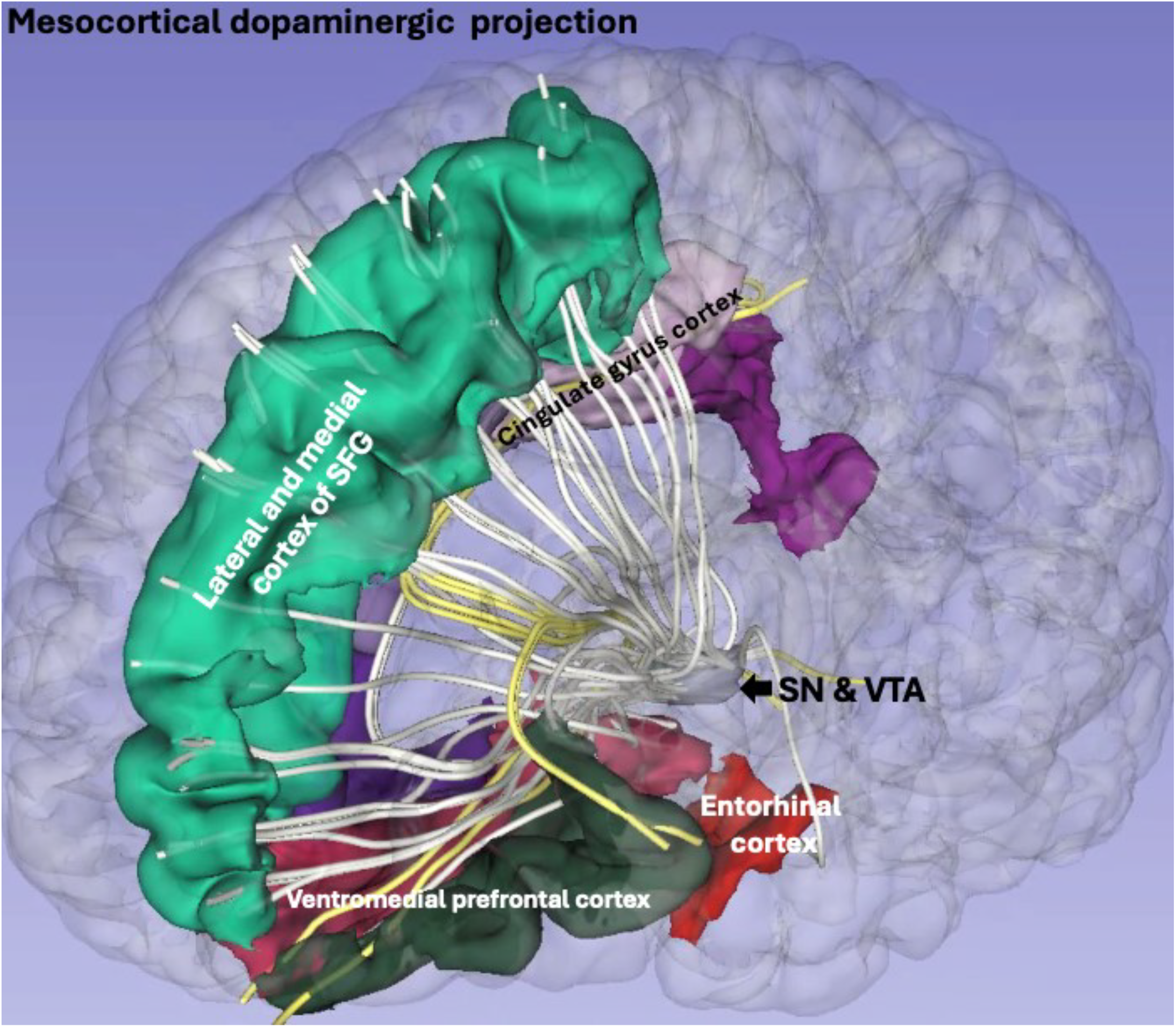
Mesocortical dopaminergic (DA) projection system involving connections from the substantia nigra (SN) and the ventral tegmental area (VTA) to the cingulate gyrus, entorhinal cortex, ventromedial prefrontal cortex and the superior frontal gyrus (SFG).

Overall, our preliminary results on the DA circuits are a groundbreaking proof of principle demonstrating that currently used multispectral clinical neuroimaging allows the detection and delineation of these neurochemical pathways. The sequence of **Figures 4, 6, 7 and 8** shows eleven individual connections of this circuitry (**Figure 4**) as well as the assembly of individual fiber connections into the three classical projectional subsystems of the mesotelencephalic DA system, namely the mesostriatal (**Figure 6**), mesolimbic (**Figure 7**) and mesocortical (**Figure 8**) projections. We expect these findings to guide further anatomical investigations in basic and clinical neuroscience. Thus, the results of this study provide a) preliminary evidence of the human DA circuits and b) strong evidence of the coerulean NE circuitry in the human brain. Furthermore, the use of a balanced sample of 12 living human subjects of the HCP repository (six females and six males matched for age) provides the basis for a preliminary normative database for the human coerulean NE circuitry.

### 3.1. Quantitative Analysis

The presence or absence of specific fiber tracts in the left or right hemisphere of each individual subject was recorded and the findings were as follows. With respect to the DA connections, out of the 110 fiber tracts (22 per subject for 5 subjects), 103 (94%) were present. The seven fiber tracts that were absent included the VTA connection with the nucleus accumbens (three in the left hemisphere and one in the right), the VTA connection with the cingulate gyrus (two in the left hemisphere), and the SN connection with the putamen (one in the right hemisphere). As shown in **Table 2**, all connections of the coerulean (LC) NE circuitry were present in all subjects. These results provide preliminary evidence for the capability of current multispectral clinical neuroimaging to delineate the dopaminergic circuitry and strong evidence for generating a pilot normative database of the coerulean NE circuitry. Biophysical parameters of average fractional anisotropy (FA), axial diffusivity (AD) and radial diffusivity (RD) of the fiber tracts are reported in **Tables 3A and 3B**. As shown in **Table 3A**, 22 connections (11 in the left hemisphere and 11 in the right hemisphere) were DA-related, whereas, as shown in **Table 3B**, four (two in the left hemisphere and two in the right hemisphere) were coerulean (LC) NE-related. Specifically, the parameters of the DA connections in five healthy human subjects are reported in **Table 3A**. Likewise, the parameters of the coerulean (LC) NE connections in 12 healthy human subjects are shown in **Table 3B**. It should be noted that the present biophysical results of the coerulean NE connections, to our knowledge, are reported here for the first time.

**Table 2.** Presence of coerulean NE circuitry (NTS/DMN – LC, and LC – PVN/Hypothalamus noradrenergic pathways) in the left and right hemispheres of 12 healthy human HCP subjects matched for gender and age (identification numbers, IDs, specified in leftmost column). X denotes the presence of a specific connection. (Abbreviations: NTS, nucleus of the solitary tract; DMN, dorsal motor nucleus of the vagus nerve; LC, locus coeruleus; PVN, paraventricular nucleus of the hypothalamus)

| ID | (NTS/DMN) ROI – (LC) ROI |  | (LC) ROI – (PVN/Hypothalamus) ROI |  |
| --- | --- | --- | --- | --- |
|  | Right | Left | Right | Left |
| 100307 | X | X | X | X |
| 101107 | X | X | X | X |
| 101915 | X | X | X | X |
| 103111 | X | X | X | X |
| 103414 | X | X | X | X |
| 110411 | X | X | X | X |
| 111312 | X | X | X | X |
| 114419 | X | X | X | X |
| 118528 | X | X | X | X |
| 118932 | X | X | X | X |
| 124422 | X | X | X | X |
| 129028 | X | X | X | X |

**Table 3A.** Average FA (fractional anisotropy), AD (axial diffusivity) and RD (radial diffusivity) of the fiber tracts representing the dopaminergic (DA) system connectivity in five healthy human subjects of the publicly available HCP repository.

| Connection | FA |  | AD |  | RD |  |
| --- | --- | --- | --- | --- | --- | --- |
|  | Right | Left | Right | Left | Right | Left |
| SN (A9 cell group) – Thalamus | 0.62576 | 0.68158 | 0.00201 | 0.0022 | 0.00049 | 0.00057 |
| SN (A9 cell group) – Putamen | 0.64402 | 0.65731 | 0.0021 | 0.00182 | 0.00057 | 0.00049 |
| SN (A9 cell group) – Entorhinal | 0.45068 | 0.46522 | 0.0017 | 0.00182 | 0.0006 | 0.00065 |
| SN (A9 cell group) – Superior Frontal | 0.70697 | 0.70201 | 0.00226 | 0.00187 | 0.00044 | 0.00044 |
| SN (A9 cell group) – Caudate | 0.69667 | 0.6306 | 0.00186 | 0.00178 | 0.00053 | 0.00051 |
| SN (A9 cell group) –<br>Cingulate | 0.58114 | 0.56618 | 0.00191 | 0.00183 | 0.00047 | 0.00047 |
| SN (A9 cell group) –<br>Orbitofrontal | 0.62306 | 0.62081 | 0.00179 | 0.00189 | 0.00053 | 0.00066 |
| VTA (A10 cell group) –<br>Thalamus | 0.70915 | 0.5457 | 0.0019 | 0.00183 | 0.00046 | 0.11621 |
| VTA (A10 cell group) –<br>Nucleus Accumbens | 0.61107 | 0.42017 | 0.00216 | 0.00151 | 0.0007 | 0.00045 |
| VTA (A10 cell group) –<br>Cingulate | 0.49872 | 0.45379 | 0.00181 | 0.00158 | 0.00059 | 0.00034 |
| VTA (A10 cell group) –<br>Orbitofrontal | 0.49678 | 0.48591 | 0.00179 | 0.00177 | 0.00058 | 0.00058 |

**Table 3B.**
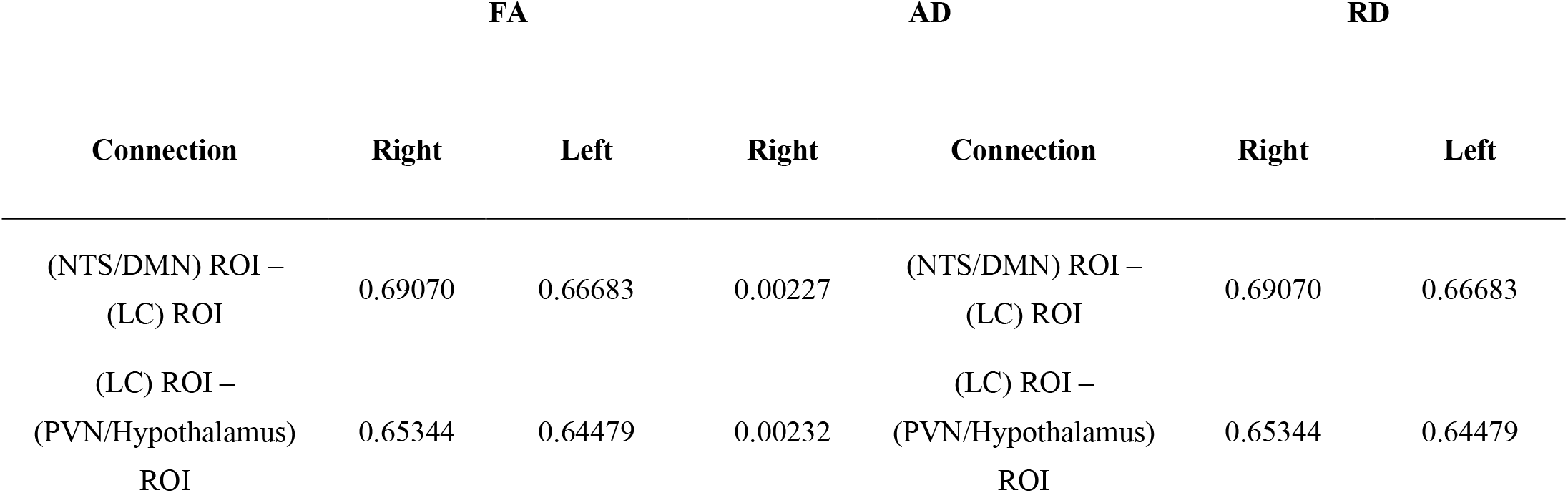
Average FA (fractional anisotropy), AD (axial diffusivity) and RD (radial diffusivity) of the fiber tracts representing structural connections of the locus coeruleus (LC) NE system in five healthy human subjects of the publicly available HCP repository. This is an original tabulation based on data from Makris et al. (6).

## 4. Discussion

In the present study, we report for the first time initial pilot results on the human dopaminergic (DA) circuitry and a preliminary normative database of coerulean noradrenergic (NE) circuitry using a multispectral imaging methodology. In a prior study we presented preliminary evidence on the human coerulean norepinephrine (NE) circuitry (6) in a small subject sample. Importantly, in the present study, by increasing the sample by a factor of more than two, with balance of age and gender, we initiated a normative database for the human coerulean NE connectivity. Moreover, to our knowledge, this is the first time that structural circuitry of the three DA systems, i.e., mesostriatal, mesolimbic and mesocortical, has been delineated in the human brain using multispectral neuroimaging, specifically combining T1-weighted MRI morphometric analysis and dMRI tractography. Although preliminary, our findings demonstrate the capability of current clinical multispectral neuroimaging to delineate the DA circuitry that is critically involved in several biobehaviors and clinical conditions such as Parkinson’s disease (PD) and schizophrenia.

### 4.1. Structural anatomy and topographical organization of catecholamine systems

The central catecholaminergic systems are associated with DA, NE and E, which constitute the catecholamine neurotransmitters in the mammalian brain. These chemical systems are organized by means of a) specific cells of origin, where these molecules are produced, b) fields or areas of termination, where the axons of the neurons producing these molecules terminate, and c) specific fiber pathways of which their axons are a part as they course from their neurons of origin to their target neurons of destination. Below, we present an account of the neuroanatomy of these chemical neural systems. It should be appreciated that the classical structural connectivity of long fiber connections is grounded principally in experimental animal studies, thus the narrative that follows has been based mainly on the animal literature (3,15,23,30).

#### DA circuitry

Dopamine (DA) neurons of origin and DA production are located mainly in the brainstem and hypothalamus. Cell hubs producing DA in the cerebrum (or telencephalon) are found in the olfactory system and the outer zone of the olfactory bulb in particular, which is referred to as the A16 cell group (31,32). The main sites of DA neurons are the mesencephalon (midbrain) and the diencephalon. Specifically, three groups of DA cells are located in the midbrain, namely cell group A8 in the retrorubral area (i.e., dorsal to the red nucleus) of the lateral tegmentum (lateral tegmental area, LTA) (e.g., (33)); cell group A9, which is formed by the pars compacta of the substantia nigra (SNpc); and cell group A10, aggregated around the midline of the ventral tegmentum and referred to as ventral tegmental area (VTA) (22–24,34). The latter is situated medial to the SN and ventral to the red nucleus. In the present MRI analysis we were not able to subdivide the SN into pars compacta and pars reticulata, thus we sampled it as a single ROI. Diencephalic structures of origin for DA are located principally in the hypothalamus, namely cell groups A12, A14 and the paraventricular nucleus (PVN) as well as in the zona incerta (ZI) and its vicinity, more specifically cell groups A11 and A13. The connections of the DA system follow a medial (or “meso”) trajectory within the neuraxis and extend within and between different subdivisions of the CNS. Given its topography and structural connectivity, the DA system was originally labeled the “mesotelencephalic dopaminergic system” referring mainly to the entire DA forebrain projection of the midbrain cell groups of its origin (23,35,36). This massive ascending mesencephalic projection, coursing principally from cell groups A8 (LTA), A9 (SNpc) and A10 (VTA), consists of two principal systems that were named by Ungerstedt (1971) as the “nigrostriatal system” and the “mesolimbic system”. Eventually, Niewenhuys (1985) proposed a more practical typology by subdividing the DA system into three principal projections and seven additional ancillary connections as follows: a) the “mesostriatal projection”, coursing from the LTA (A8 cell group), SNpc (A9 cell group) and VTA (A10 cell group) (e.g., (23,36,37); b) the “mesolimbic projection”, originating from the VTA (A10 cell group) (3); and c) the “mesocortical projection”, which originates from the VTA (A10 cell group) and the medial portion of the substantia nigra. The mesostriatal projection connects the SNpc (A9 cell group) principally with the caudate nucleus and putamen. It also connects the VTA (A10 cell group) mainly with the nucleus accumbens septi (NAc), and the LTA (A8 cell group) with the ventral putamen (3,38–40). The mesolimbic projection connects the VTA with the NAc and ascends within the medial forebrain bundle (MFB), in a location medial to the mesostriatal projection. It also connects the VTA with olfactory areas such as the olfactory bulb and tubercle and the anterior olfactory nucleus. Furthermore, it connects the VTA with the anterior perforated substance, the lateral septal nucleus, the bed nucleus of the stria terminalis and, importantly, with the central and basal nuclei of the amygdala. The mesocortical projection originates from the VTA and the medial portion of the substantia nigra, connecting with the medial aspect of the frontal lobe in each cerebral hemisphere, namely the piriform and pre-piriform olfactory cortices, the entorhinal cortex and, notably, the anterior cingulate cortex (ACC) (3,38–40).

In addition to the three major DA projections described above, there are four efferent hypothalamic and three mesencephalic ancillary projections in the dopaminergic system. Hypothalamic DA projections arise from four distinct regions as follows: a) The tubero-infundibular area cell group A12, which projects to the median eminence and the neurohypophysis (i.e., the posterior pituitary gland) (e.g., (3)); b) the A11 and A13 cell groups in the region of the zona incerta, which form local connections within adjacent hypothalamic nuclei and are termed the incerto-hypothalamic projection (e.g., (3)); c) the A11, A13 and A14 cell groups, which project to the lateral septal nucleus and constitute the diencephalic-septal dopaminergic projection (41); and d) the PVN and the A11 and A13 DA cell groups of the ZI, which project to the spinal cord giving rise to the hypothalamo-spinal dopaminergic projection. Furthermore, Swanson and colleagues (1981) (42) reported a projection from the PVN to the dorsal vagal complex (DVC, comprising the nucleus of solitary tract or NTS and the dorsal motor nucleus of the vagus or DMN). It is notable that the hypothalamo-spinal dopaminergic projection courses via the dorsal periventricular catecholaminergic bundle, which is part of the dorsal longitudinal fasciculus of Schutz (DLF) (36,43). Moreover, three additional ancillary efferent DA projections of mesencephalic origin have been reported as follows: a) a projection from the VTA to the lateral habenular nucleus via the fasciculus retroflexus; b) a projection from the SNpc to the neighboring subthalamic nucleus (STN); and c) a bilateral projection from VTA to the locus coeruleus (e.g., (3)).

#### Coerulean NE circuitry

Norepinephrine (NE) neurons of origin are encountered principally in the brainstem. The coerulean NE circuitry comprises extensive connections of LC cells of origin with the cerebrum and spinal cord. The fiber tracts of coerulean origin fasciculate course within the dorsal noradrenergic bundle [e.g., 40,44) and the dorsal longitudinal fasciculus of Schutz (3,44). En route to the diencephalon and telencephalon, these fiber pathways contribute to the medial forebrain bundle (MFB) [e.g., 42,44). Thus, the LC is linked with brain structures such as the PVN rostrally and the NTS and DMN caudally. Finally, there is a non-coerulean NE circuitry also arising from the brainstem. Together with the pontine coerulean NE complex, constituted principally by cell group A6/LC and by adjoining noradrenergic neurons in the central gray and subcoerulean regions, they comprise the central NE brain system. The non-coerulean cell groups of origin of the NE system, such as the A2 cell group in the upper (or dorsal) medulla and the A1, A5 and A7 cell groups in the pontine lateral tegmentum, share with the coerulean complex approximately a 50/50 contribution to the origin of the NE central system. They jointly constitute a long ascending fiber tract called the ventral noradrenergic pathway or bundle (VNP or VNB), which courses mainly within the MFB (45).

### 4.2. Functional and clinical roles of central catecholamine systems

Depending on the specific circuit, <u>the dopaminergic (DA) system</u> manifests different effects. The mesostriatal DA projection provides the ability to switch motor programs (46–48), which is a highly valuable behavior for a biological organism to operate promptly and efficiently under constantly changing environmental scenarios, especially in the natural world where motor functions are critical for predatory and defensive behaviors. This ability to flexibly switch motor programs extends to cognitive behaviors as well, which is especially relevant in humans. Dopamine works in balance with other neurotransmitters during movement. In particular, it seems that dopamine coordinates the voluntary, or pyramidal, and involuntary, or extrapyramidal, systems. Dopamine deficit is considered to be the principal cause of Parkinson’s disease, a movement disorder characterized by uncontrolled movements, stiffness, slowing of movement and difficulty with balance that raises the risk of falls. Changes in dopamine levels have been associated with conditions other than Parkinson’s disease, such as restless legs syndrome and attention deficit hyperactivity disorder (ADHD). The mesolimbic DA projection of the dopaminergic system plays a role in the experience of pleasure and reward as well as in many other functions, including, e.g., memory, movement, motivation, mood, and attention. This is reflected via the connections of SNpc with the NAc, which are strongly related to reward functions and are considered as the principal “reward center” in the brain (e.g., (49–51)). The reward network may be an integral part of the neurobiology of drug addiction (52). Structural alterations in this system may enhance the risk for reward deficiency syndrome (53,54) and drug-seeking behaviors. This circuitry seems to be critical to abnormalities in hedonic set points associated with increased drug use and drug dependence (52,55). By contrast, when there is a decrease of dopamine, depression-like symptoms may appear, such as apathy or feelings of hopelessness. Furthermore, hyperactivity of the VTA and its dopaminergic outflow to limbic structures may play a role in schizophrenia (56–59).

<u>The noradrenergic (NE) system</u> is related to a range of functions depending on the specific brainstem cell groups and their associated circuits. In general, the LC complex innervates sensory and associative nuclei and brain centers, whereas the non-coerulean NE cell groups are related to visceral and motor functions. More specifically, the LC NE complex has been shown to play a critical role in attention by acting as an “alarm-like” system that monitors the environment and prepares the organism to face putative or real emergency situations (e.g., (6)). These reactions can elicit increases in heart rate and blood pressure similar to “fight-or-flight” responses. Furthermore, prolonged negative stress can produce lasting structural and physiological changes in the LC and “DVC-LC-PVN circuitry”, a condition that may increase the risk not only for cardiovascular disease but also for major depressive disorder (MDD) and posttraumatic stress disorder (PTSD) (e.g., (6)). Furthermore, this circuitry is essential for the preservation of cognitive and affective functions, which is critically important for the aging brain (60). By contrast, the non-coerulean NE system is involved in a variety of viscero-effector and neuroendocrine functions for cardiovascular control and respiration. In the cardiovascular domain, these responses result in tachycardia and increased blood pressure. Specifically, the NE circuitry including the locus coeruleus and its connections with the NTS and DMN (which comprise the dorsal vagal complex or DVC) constitutes an autonomic pathway that stimulates the paraventricular nucleus of the hypothalamus (PVN) to secrete vasopressin or antidiuretic hormone (ADH) (eg, (61,62). This circuitry is critically relevant in arterial blood pressure regulation and cardiac function; consequently, their dysfunction could well lead to cardiovascular disease (eg, (1,3,63–68). Finally, the adrenergic system is associated mainly with blood pressure regulation (e.g., (3)).

### 4.3. Brain-immune interactions and clinical implications of catecholaminergic systems for COVID-19

Brain-immune interactions involve a reflexive hard-wired circuit of the nervous system (NS) (69,70), which is characterized by inflammation-sensing and inflammation-suppressing functions (26). Modulation of immunity by the NS has been highly informative with respect to the anatomical and molecular mechanisms underlying the brain-immune dynamics during inflammation. The afferent component of the vagus nerve can be stimulated by pro-inflammatory cytokines produced by the peripheral inflammatory process in SARS-CoV-2 virus infection (25,26,71,72). This information is conveyed to the NTS in the medulla and to the NTS-DMN ensemble (or DVC) where the coordination of vagal afferent and efferent responses takes place (25,26,73); thus the vagus is able to suppress inflammation in real time (26,73). Although the complete neural circuitry involved in immune modulation remains to be identified with clarity, it seems plausible that the catecholaminergic system, via the LC and propriobulbar connections of the non-coerulean NE groups as well as the adrenergic cell groups with the DVC, plays a key modulatory role in this regard. It should be noted that the neuroimmune circuitry could constitute a more extensive network involving neural centers and fiber pathway connections across the CNS, including the cerebral cortex, subcortical nuclei, the limbic and paralimbic systems as well as the cerebellum and brainstem. It is expected that multispectral neuroimaging will be a highly valuable asset in this regard, given its non-invasive and in vivo nature. This capability will open a number of novel avenues in multispectral neuroimaging research for traditional and chemical neuroanatomy as well as several clinical opportunities for bridging neuroimaging with histopathology as discussed in detail in a recent publication by our group in which we elaborate upon the particular case of acute and post-acute (or long) COVID-19 (74).

### 4.4. Limitations and future studies

Brainstem neuroanatomical investigations using neuroimaging present several limitations. T1- and T2-weighted MRI-based morphometric approaches for human subject in vivo investigations, as used in the present study, lack the necessary spatial resolution and signal-to-noise ratio to make them comparable to actual neuroanatomical and histopathological observations. This is especially the case in brain regions such as the substantia nigra and several other structures in the brainstem, which are characterized by highly dense, heterogeneous and complex anatomical architecture. Likewise, dMRI tractography studies can be only approximate compared to the tract tracing observations derived from experimental animal studies. Thus, we addressed these limitations by using an ultra-high-resolution human dataset of the brainstem and diencephalon, which allowed us to approximate significantly the topography and connectivity of these circuitries to what is known from the experimental animal connectivity literature (e.g., (2)). Notably, our knowledge of structural connectivity of the human brain has been derived principally from other animals, such as rodents, cats, dogs or nonhuman primates as has been elaborated upon in previous reports by our group and others (e.g., (15)). Taking into account these key considerations we investigated only aspects of the human central catecholaminergic structural systems associated with structures that we identified reliably in our studies of post-mortem and in vivo MRI human datasets. These structures included the hypothalamic subregions (12) as well as the locus coeruleus, substantia nigra and VTA (14), which we have studied in detail using combined histology, ultra-high-resolution and clinical low-spatial-resolution hypothalamic MRI (12) and combined ultra-high-resolution and clinical low-spatial-resolution brain-stem MRI (14). Herein we demonstrated that it is possible to delineate several fiber tracts associated with the central catecholaminergic systems. Notably, we were able to demonstrate reliably the coerulean NE connections, i.e., the NTS/DMN – LC and the LC – PVN/Hypothalamus noradrenergic pathways in all subjects, using current multispectral clinical imaging techniques. With respect to the DA connections, the results were slightly less consistent, i.e., 94% of the DA pathways were observed across subjects compared to 100% of the NE pathways (as elaborated upon in the results). Inconsistency was encountered in the VTA connection with the nucleus accumbens, the VTA connection with the cingulate gyrus, and the SN connection with the putamen. We believe that the difficulty in delineating these three DA connections may be related to the complex architecture of the white matter between the VTA/SN brainstem region and the cerebral cingulate, caudate nucleus and putamen regions. This difficulty could be attributed to the massive number of white matter fibers, e.g., of the anterior limb of the internal capsule and the corticospinal tract, that connect the cerebrum with the brainstem and spinal cord. This issue did not arise with respect to the connections of the coerulean NE pathways given that the level of white matter complexity is not as pronounced in the dorsal brainstem. In conclusion, although methodologically challenging and to a certain degree approximate, this study has involved a great deal of sophistication in both the neuroanatomical domain and in multispectral neuroimaging analysis. Our current findings for the coerulean NE system and our preliminary results for the DA system are highly encouraging for studying these structures clinically using MRI. Moreover, our results clearly indicate that for clinical applications, further technological advances are needed to produce neuroimaging protocols with optimal voxel size and increased signal-to-noise ratios. Furthermore, future functional and metabolic investigations using fMRI and PET, respectively, could enable us to advance diagnosis and monitoring of treatment targeting the brain-immune related central catecholamine circuitries in medical research and practice.

## 5. Conclusions

Overall, the results of the present study approximate substantially the known structural neuroanatomy of the human noradrenergic and dopaminergic circuitries as described in the experimental animal literature. Current multispectral neuroimaging allows the study of clinical conditions associated with the central catecholamine circuitries such as cardiovascular disease, major depression, schizophrenia, and other disorders associated with chronic stress. Furthermore, LC as well as propriobulbar connections (3) link the NE cell groups with several nuclei such as the NTS and DMN that constitute the DVC (3,75,76). These specific connections seem especially relevant for brain-immune interactions and for clinical implications of the central catecholaminergic systems in diseases such as SARS-CoV-2 virus infection. Notably, the current advances in neuroimaging techniques allow us to identify and assess clinically or in post-mortem settings specific brain circuits such as those associated with the neuroimmune network. This capability will empower basic and clinical research in an unprecedented way, especially due to its applicability for facing dire medical challenges such as acute COVID-19 and post-acute or long COVID. Importantly, we were able to delineate the mesostriatal, mesolimbic and mesocortical DA projection systems and to reconstruct them in three-dimensional space. This was achieved in routine MRI datasets of HCP healthy human subjects, which indicates applicability to clinical populations using neuroimaging. By elucidating the coerulean NE and DA central catecholamine circuitries in the human brain using multispectral neuroimaging we demonstrated the great potential of translating such difficult and complex anatomical information from experimental animals to humans. We expect this pioneering neuroanatomical research to encourage further technological development in neuroimaging that will empower clinical applications with the use of MRI protocols allowing higher signal-to-noise ratios and higher spatial resolution.

## 6. Patents

Not Applicable.

### Supplementary Materials

Not Applicable.

### Author Contributions

All authors have read and agreed to the published version of the manuscript. Conceptualization: Nikos Makris and Agustin Castañeyra-Perdomo; Investigation: Nikos Makris, Poliana Hartung Toppa, Richard J. Rushmore, Kayley Haggerty, George Papadimitriou, Agustin Castañeyra-Perdomo; Methodology: All authors; Writing – original draft: All authors; Writing – review & editing: All authors.

### Funding

We would like to acknowledge the following grants for their support: R01MH112748 (NM, RJR), R01AG042512 (NM), K24MH116366, R01MH132610, R01MH125860 (NM), and R21NS136960 (RJR, NM), R01NS125307 (RJR, NM).

### Institutional Review Board Statement

Not Applicable.

### Informed Consent Statement

Not applicable.

### Data Availability Statement

Data were provided in part by the HCP, WU-Minn Consortium (Principal Investigators: David Van Essen and Kamil Ugurbil; 1U54MH091657) funded by the 16 NIH Institutes and Centers that support the NIH Blueprint for Neuroscience Research, and by the McDonnell Center for Systems Neuroscience at Washington University. https://www.humanconnectome.org/study/hcp-young-adult/document/hcp-citations

## Acknowledgments

We would also like to recognize Drs. Allan Johnson and Evan Calabrese for providing excellent post-mortem data to the neuroscience community, in particular the brainstem/diencephalon human dataset used in this study.

## Appendix A

Not Applicable.

## Appendix B

Not Applicable.

## Disclaimer/Publisher’s Note

The statements, opinions and data contained in all publications are solely those of the individual author(s) and contributor(s) and not of MDPI and/or the editor(s). MDPI and/or the editor(s) disclaim responsibility for any injury to people or property resulting from any ideas, methods, instructions or products referred to in the content.

